# The fern CYPome: Fern-specific cytochrome P450 family involved in convergent evolution of chemical defense

**DOI:** 10.1101/2021.03.23.436569

**Authors:** Sara Thodberg, Cecilie Cetti Hansen, Adam M. Takos, Martina Pičmanová, Birger Lindberg Møller, David R. Nelson, Elizabeth H. Jakobsen Neilson

**Affiliations:** Plant Biochemistry Laboratory, Department of Plant and Environmental Sciences, University of Copenhagen, Thorvaldsensvej 40, 1871 Frederiksberg C, Denmark; SNIPR Biome ApS, Lersø Parkallé 44, 2100 Østerbro, Denmark; Institute of Quantitative Biology, Biochemistry and Biotechnology, University of Edinburgh, C.H. Waddington Building, Max Born Crescent, Edinburgh EH9 3BF, United Kingdom; Department of Microbiology, Immunology and Biochemistry, University of Tennessee, 858 Madison Ave. Suite G01, Memphis TN 38163

## Abstract

Plant natural products encompass an enormous chemical diversity bearing witness to great molecular innovation that occured throughout land plant evolution. Cytochrome P450 monooxygenases (CYPs) catalyze a wide variety of monooxygenation reactions essential to the metabolic repertoire of plants natural products. Ferns constitute the second largest group of vascular plants and hold a significant phylogenetic position in land evolution, lying sister to seed plants. To date, CYP diversity has not been described for this taxon and pathway discovery in ferns in general is scarce, despite possessing a rich diversity of natural products. We analysed over 8000 available fern CYPs, classifing and characterizing the landscape of this super-enzyme group. Fern CYPs are dominated by fern-specific families (∼60%), with the largest family – CYP981 – constituting approximately 15% of all predicted fern CYPs in the dataset. The abundancy and dynamics of the CYP981 family suggest a position equivalent to the CYP71 family present in seed plants, with potential roles in natural product biosynthesis. Ferns are the evolutionary oldest group to biosynthesize cyanogenic glycosides; amino acid-derived defense compounds. We show that CYP981F5 from the highly cyanogenic fern *Phlebodium aureum* catalyzes the conversion of phenylacetonitrile to mandelonitrile, an intermediate step in cyanogenic glycoside biosynthesis. The fern CYPome provides an important platform to further understand evolution of metabolite biosynthesis throughout the plant kingdom, and in ferns specifically.

## Introduction

Plants biosynthesize an enormous diversity of bioactive natural products to interact with their biotic and abiotic environment. These natural products (also known as specialized metabolites) are involved in an immense number of roles including defense against herbivory and pathogen attack, attraction of pollinators, signaling or as protective agents against abiotic stress factors. Some natural products classes, such as the phenylpropanoids, are conserved throughout many plant lineages. Other classes, such as glucosinolates and benzoxazinoids, are constrained to certain phylogenetic groups demonstrating an acquired ability to biosynthesize niche-specific natural products as a result of evolutionary adaptions to environmental challenges. New pathways can arise via gene duplication and neofunctionalization and advantageous adaptions are maintained through positive selection (Feyereisen, 2011; Weng, 2014).

Plant cytochromes P450 (CYPs) catalyze a wide variety of coveted monooxygenation reactions in both general and specialized metabolism expanding the metabolic repertoire of plants. Diversification within the CYPome (the total number of CYP genes in a given species or taxonomic group) has led to the emergence of new metabolic pathways playing pivotal roles in land plant evolution (Nelson and Werck-Reichhart, 2011). Within recent years, genome and transcriptome sequencing efforts have resulted in an explosion in the number of reported gene sequences encoding CYPs. The large number of CYPs are sorted into families and subfamilies based on >40% and >55% amino acid sequence identities and by phylogenetic grouping (Nelson et al., 1996; Nelson, 2009). The plant CYP families hold the numbers CYP51, 71-99, 701-999 and from 7001-9999 (Nelson, 2018).

Ferns (class Polypodiopsida), are a very diverse class of vascular plants consisting of approximately 10,600 different species (I P P G, 2016). Ferns are comprised of four sub-classes – Equisetidae (horsetails); Ophioglossidae; Marattiidae; and Polypodiidae (leptosporangiates) – of which the Polypodiidae constitute ∼94% of all fern species. Together, ferns represent the second largest group of plants following angiosperms, with the major derived lineages within the Polypodiidae (primarily the order Polypodiales), diversifying following the expansion of angiosperms during the late Cretaceous period approximately 100 Mya (Schneider et al., 2004). From an evolutionary perspective, ferns hold an important position in plant evolution, lying in the sister clade to seed plants. Unlike seed plants, however, ferns do not undergo the typical genome downsizing following whole genome duplication events (Liu et al., 2019). This has resulted in ferns possessing relatively higher chromosome numbers and genome sizes compared with angiosperms (Sessa et al.; Wolf et al., 2015; Clark et al., 2016; Li et al., 2018)). This genome complexity has meant that only two fern species have been fully sequenced to date (Li et al., 2018), and the fern CYPome has not yet been described.

Ferns produce a large variety of toxic, anti-nutritional and oncogenic natural products including cyanogenic glycosides, ecdysones, tannins and ptaquilosides (Alonso-Amelot, 2002; Popper and Fry, 2004; Kisielius et al., 2020), with their biosynthesis presumably involving CYPs. The aforementioned large and complex genomes of ferns have also hindered genome-guided elucidation of natural product biosynthetic pathways. Advancement of third generation sequencing and initiatives such as the OneKP database (Auger et al., 2014; Matasci et al., 2014; Leebens-Mack et al., 2019) provides a critical resource for the identification and characterization of biosynthetic genes from otherwise less-studied and genetically complex species within the fern class.

Ferns are the oldest evolutionary plant group known to synthesize cyanogenic glycosides (Bak et al., 2006), specifically the phenylalanine-derived mono-glucoside prunasin, and di-glycoside vicianin (Schreiner et al., 1984; Lizotte et al., 1986). As CYPs are typically involved in cyanogenic glycoside biosynthesis, this class of natural product provides an excellent system by which to undertake pathway elucidation and the characterization of CYPs in ferns (Møller, 2010; Thodberg et al., 2020). Cyanogenic glycosides are present in more than 3000 plant species providing chemical defense against herbivory, in a process termed cyanogenesis (Conn, 1980; Poulton, 1990; Gleadow and Møller, 2014a). This two-component defense system is detonated upon tissue disruption, causing β-glucosidase-mediated hydrolysis of the stable glucosylated α-hydroxynitrile and the resultant release of toxic hydrogen cyanide gas (Gleadow and Woodrow, 2002; Morant et al., 2008). In addition to chemical defense, cyanogenic glucosides have also been shown to play an important roles in general metabolism such as providing a storage form of reduced nitrogen, and in controlling flowering time (Møller, 2010; Neilson et al., 2011; Pičmanová et al., 2015; Del Cueto et al., 2017; Ionescu et al., 2017b; Ionescu et al., 2017a; Bjarnholt et al., 2018; Schmidt et al., 2018).

Cyanogenic glycosides are derived from the amino acids phenylalanine, tyrosine, valine, leucine, and isoleucine, and from the non-protein amino acid cyclopentenylglycine (Olafsdottir et al., 1991; Gleadow and Møller, 2014). The full biosynthetic pathway for cyanogenic glucosides has been characterized in several angiosperm species including sorghum (*Sorghum bicolor;* (Koch et al., 1995; Bak et al., 1998b; Jones et al., 1999), cassava (*Manihot esculenta;* Andersen et al., 2000; Jørgensen et al., 2011; Kannangara et al., 2011), barley (*Hordeum vulgare;* Knoch *et al*., 2016), *Lotus japonicus* (Forslund, 2004; Takos et al., 2011), *Eucalyptus cladocalyx* (Hansen et al., 2018), almond (*Prunus dulcis*; Thodberg et al., 2018) and green bean (*Phaseolus vulgaris;* Lai et al., 2020), and has been partially identified in conifers (Luck et al., 2017). Comparison of the biosynthetic pathways reveal that the biosynthesis has arisen independently several times in the plant kingdom (Takos et al., 2011; Hansen et al., 2018; Thodberg et al., 2020).

The first step of cyanogenic glucoside biosynthesis in gymnosperms and angiosperms is functionally conserved and catalyzed by a CYP79, which converts the precursor amino acid into a corresponding oxime (Luck et al., 2017; Sørensen et al., 2017). We recently showed that ferns do not contain CYP members of the CYP79 family but instead possess a flavin monooxygenase, termed FOS1, that catalyzes the conversion of phenylalanine to phenylacetaldoxime (Thodberg et al., 2020). In angiosperms, conversion of the oxime to the corresponding cyanohydrin is typically catalyzed by a multifunctional CYP from the CYP71 or CYP736 family (Bak et al., 1998a; Takos et al., 2011). As with the CYP79 family, both CYP71s and CYP736s are absent from ferns (Nelson and Werck-Reichhart, 2011; Thodberg et al., 2020). The cyanohydrin is formed by the structural re-arrangement of the E-oxime into a Z-oxime, followed by a dehydration and C-hydroxylation (Clausen et al., 2015). In *Eucalyptus*, these reactions are catalyzed by two distinct CYPs: CYP706C55 catalyzes a dehydration reaction of the oxime to form a nitrile, which is subsequently hydroxylated by CYP71B103 forming the cyanohydrin (Hansen et al., 2018). Finally, the cyanohydrin is stabilized by glucosylation catalyzed by a UGT85 (Jones et al., 1999; Franks et al., 2008; Kannangara et al., 2011; Takos et al., 2011; Hansen et al., 2018), or also by a UGT94 in almond (Thodberg et al., 2018).

Here, we utilized the OneKP database to investigate the fern CYPome and identified a CYP-rich fern-specific family reminiscent of the CYP71 family in seed plants, designated CYP981. To assess if CYP981 members are involved in cyanogenic glycoside biosynthesis, we mined our previously published transcriptomic dataset of the highly cyanogenic species *Phlebodium aureum* (Thodberg et al., 2020). Functional characterization of the differentially expressed *CYP981F5* revealed its involvement in production of phenylalanine-derived cyanogenic glycosides, by catalyzing the hydroxylation of phenylacetonitrile to the corresponding cyanohydrin. This example of convergent evolution underlines the dynamics of CYP recruitment to specific biosynthetic pathways.

## Results

### CYPome of Ferns

To investigate the CYPome of ferns, a TBLASTN search against the 69 fern transcriptomes in the OneKP dataset was performed (Table S1). This BLAST identified approximately 8930 contigs encoding full length and fragment CYP sequences (Data S1). Annotation of the 8000+ fern-derived CYP contigs was performed according to the internationally used nomenclature system (Nelson, 2006), resulting in their assignments to 79 different CYP families. This identified ferns as having the highest number of CYP families compared to angiosperms, gymnosperms, lycophytes, hornworts, moss and liverworts (Fig. 1). Of these 79 CYP families, 49 families (60%) are exclusive to ferns (Fig.1 and Fig. 2A).

**Figure 1.**
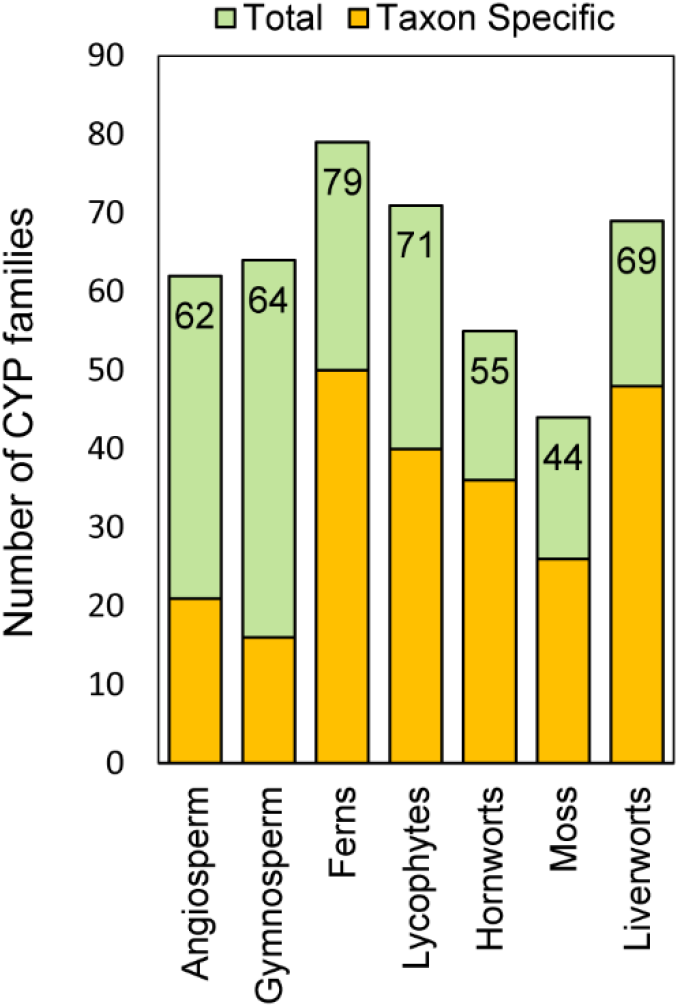
Diversification of CYP families across land plant evolution. Number of CYP families in land plant taxa. Ferns have the largest number of CYP families (79) with 49 families (60%) as novel fern specific families

**Figure 2.**
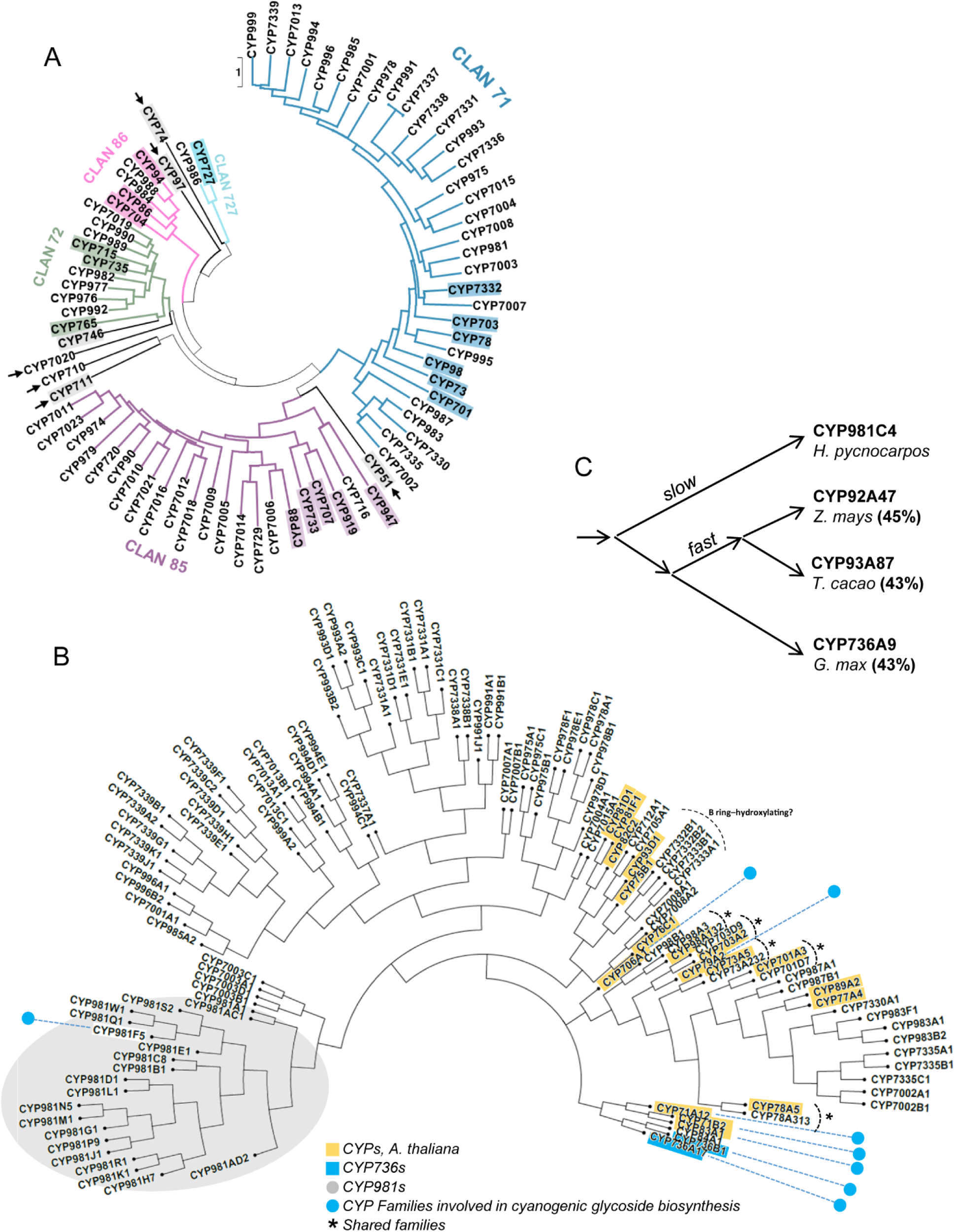
CYPome of ferns. A) Phylogenetic tree of CYP families in ferns with a total of 79 sequences, where each family is represented by a single member. Families shared with other taxa appears with colored background. The 12 CYP clans are annotated and dominant clans are highlighted according to colors. The smaller clans are marked by arrows and namesaked by the represented members. The tree is rooted to CYP74. B) All subfamilies (<55%) belonging to clan 71 from ferns (black) compared to Arabidopsis thaliana (yellow). The five families shared between fern and Arabidopsis, such as CYP73 and CYP98 are marked with an asterisk. CYP736 is not present in Arabidopsis, but two representatives are included due to this family’s involvement in cyanogenic glycoside biosynthesis. The CYPs involved in cyanogenic glycoside biosynthesis (CYP79, CYP706, CYP71, CYP736 and CYP981) are marked with blue dots. C) Schematic illustration of the evolutionary pace of cytochrome P450s resulting in different cytochrome P450 families with > 40% sequence identity.

A phylogenetic tree of all annotated fern CYPs was constructed based on their amino acid sequences, and the arising clade features were analyzed (Fig. 2A). Ferns contain the nine conserved land plant CYP clans (CYP51, CYP71, CYP72, CYP74, CYP85, CYP86, CYP97, CYP710 and CYP711), a one-family member clan 7020 only known from eight fern species and lycophytes, as well as clan 746 predominantly found in algae, but also seen in liverworts, moss and three fern species (Data S1). Clans 71, 85 and 72 are dominant in ferns, similar to their dominance in other land plant taxa (Fig 2A). In particular, the 71 clan represents more than 50% of all plant CYPs, except in hornworts where they constitute 32% of the CYPs.

A smaller clan, the 727 clan is represented by the CYP986 and CYP727 families in ferns. The occurrence of the CYP986 family is seemingly restricted to the three primitive fern families outside of the Polypodiidae sub-class, Marattiaceae, Equisetaceaea and Ophioglossaceae, whereas the CYP727 family was only identified in Osmundaceae (Order: Osmundales). Looking at the occurrence of families in the 727 clan across land plants, most are distributed in a taxon-specific manner, from moss to angiosperms, a pattern indicating an orthologue relationship of these families.

Five CYP families – CYP715, CYP720, CYP729, CYP733 and CYP735 – first appear in ferns (i.e. not present in lycophytes). For example, CYP720 and CYP729 (clan 85) are present in both gymnosperms and angiosperms, CYP733 (clan 85) has at present only been found in angiosperms in addition to ferns, while CYP7005 (clan 85) is only found in ferns and gymnosperms. Other CYP families were found to arise before the evolution of ferns, but not retained in angiosperms. For example, CYP947 (clan 85) is found specifically in lycophytes, ferns and gymnosperm, whilst CYP7020 (clan 7020) is present in lycophytes, diversified in ferns, but not present in seed plants.

By comparing CYP families within the 71 clan from ferns and Arabidopsis (*Arabidopsis thaliana*), only five families are shared (CYP73, CYP78, CYP98, CYP701, CYP703, see Fig. 2B). Some fern-specific CYP families are closely related to families known from *Arabidopsis*, while other phylogenetic branches have significantly evolved and diversified specifically in ferns. Interestingly, ferns do not possess CYPs belonging to the CYP75 family despite the reported accumulation of flavonoids in which the B-ring is 3’- and 5’-hydroxylated (Hiraoka, 1978; Ren et al., 2014).

Of the conserved and characterized CYP families, several are maintained at a low copy number. Within the 69 fern transcriptomes analyzed, we find a single CYP51 and a single CYP98 in most species. This supports their conserved gate-keeping functions whereby CYP51G is a sterol 14α-sterol demethylase (Kahn et al., 1996) and CYP98A is a *p*-coumaroyl-shikimate 3’-hydroxylase (Alber et al., 2019). Notably, four fern species from the family Ophioglossaceae (Order: Ophioflossales) contain additional CYP98 copies in subfamily B. An intriguing increase in copy number is observed for the CYP73 and CYP74 families with 654 and 490 transcripts identified, respectively (i.e average of nine CYP73 and seven CYP74 copies per species).

### Identification of a highly abundant fern-specific CYP981 family

Analysis of the fern CYPome identified a large fern-specific CYP family within the 71 clan. More than 1200 sequences from the OneKP resource, equivalent to 15% of the total fern CYP sequences available, are classified as members of this family. They share >40% sequence identity to CYPs of the CYP92, CYP93 and CYP736 families, but form a distinct phylogenetic group. This CYP family was named CYP981 (Fig. 2B-C).

The transcriptome collection from fern species within the Polypodiales order contained an average of 17 members of the CYP981 family per species, much higher compared to species in the other orders (Fig. 3). The family arose in ferns, but the sub-classes Equisetidae, Ophioglossidae and Marattiidae express a comparatively low number of CYP981s compared to the leptosporangiate ferns with Polypodiidae. For example, only two CYP981s are expressed in *Ophioglosum vulgatum* compared to up to 44 in *Cystopteris utahensis* (Polypodiales).

**Figure 3.**
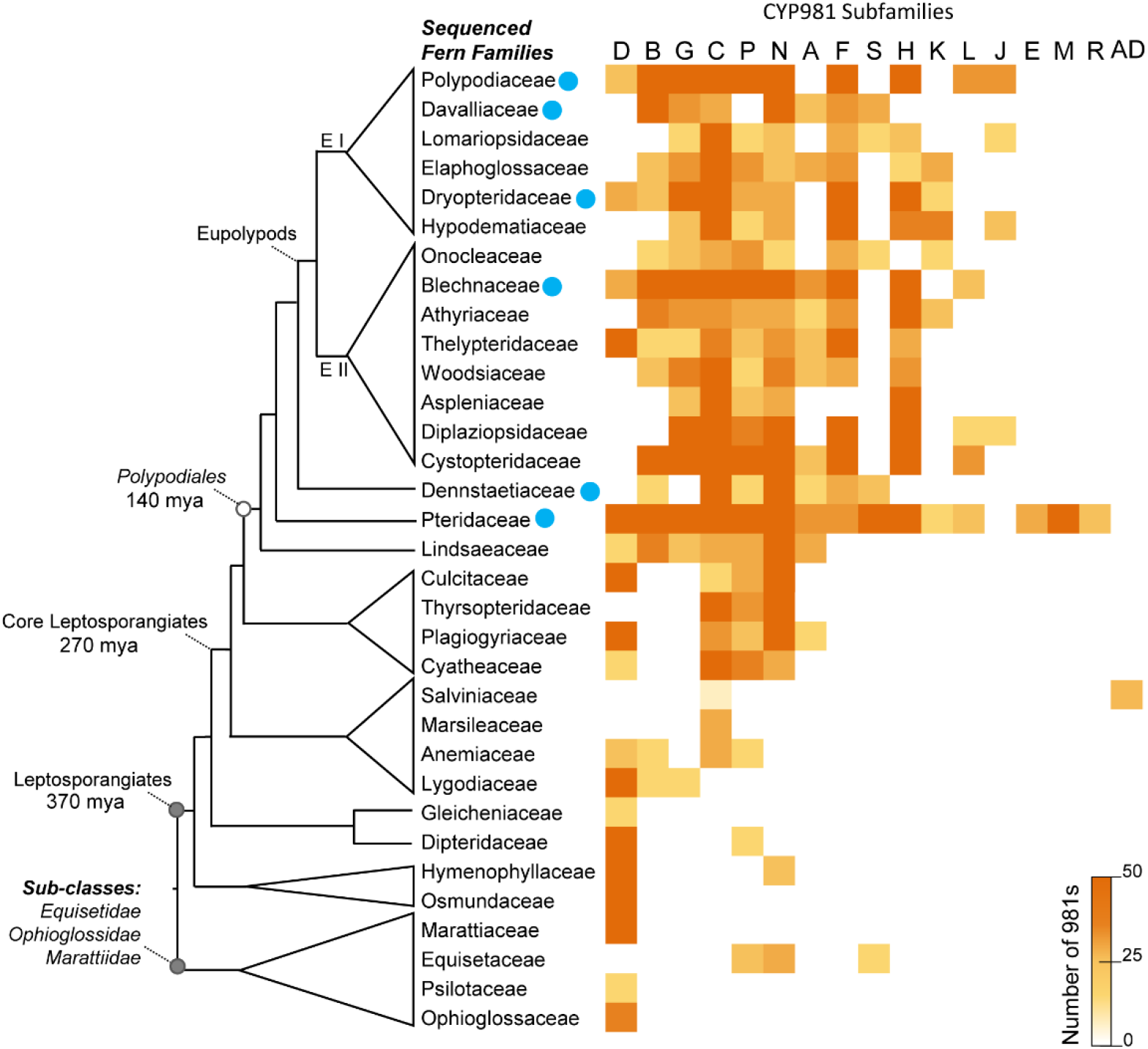
The occurrence of CYP981 subfamilies across fern evolution. Mapping of the 1200 CYP981 sequences from the OneKP database and the transcriptomes produced in the current study illustrates the evolutionary distribution and depth of this fern specific family. The fern families represented in the OneKP are grouped in broad taxonomy clades and schematically drawn according to PPGI 2016 (I P P G, 2016). Fern sub-classes (grey), orders (white) and informal clades are indicated. Blue dots denote fern families with at least one cyanogenic glucoside producing species. The CYP981 subfamilies are sorted based on their temporal appearance.

The Polypodiales exhibit broader CYP981 sequence diversity than the prior leptosporangiates as documented by the presence of a high number of CYP981 subfamilies. The subfamilies CYP981F, S, H, K, L, D, J, E, M and R appear specific to Polypodiales ferns (Fig. 3). Specifically, the subfamilies E, M and R are only found in Pteridaceae. Salviniaceae, which hold the aquatic *Azolla* fern species, have a narrow subfamily pattern, only containing subfamily C and a Salviniacea-specific subfamily AD.

Albeit the CYP981s exhibits >40% amino acid sequence identity to the CYP736 family, the CYP981s forms a distinct phylogenetic group in the clan 71 (Fig. 2C). The angiosperm CYP736A subfamily contains a few members with known functionalities, including CYP736A2 from *Lotus japonicus* which catalyzes the oxime to cyanohydrin conversion in cyanogenic glucoside biosynthesis (Takos et al., 2011). Given this high amino acid sequence identity between the CYP736 and CYP981 families, we investigated the possible involvement of CYP981 members in cyanogenic glycoside biosynthesis in ferns using the previously published comparative transcriptome of cyanogenic *P. aureum* (Thodberg et al 2020).

### CYP981F5 is involved in cyanogenic glycoside biosynthesis

Analysis of the abundant fern-specific CYP981 family in *P. aureum* identified six full-length CYP981 transcripts (Fig. 4A). Two of these, *PaCYP981F5* and *PaCYP981P7*, were highly expressed in tissue accumulating high amounts of cyanogenic glycoside (fiddlehead) and poorly expressed in tissue accumulating low amounts of cyanogenic glycoside (young pinnae). The other transcribed CYP981s showed higher expression in tissues accumulating lower concentrations of cyanogenic glycosides.

**Figure 4.**
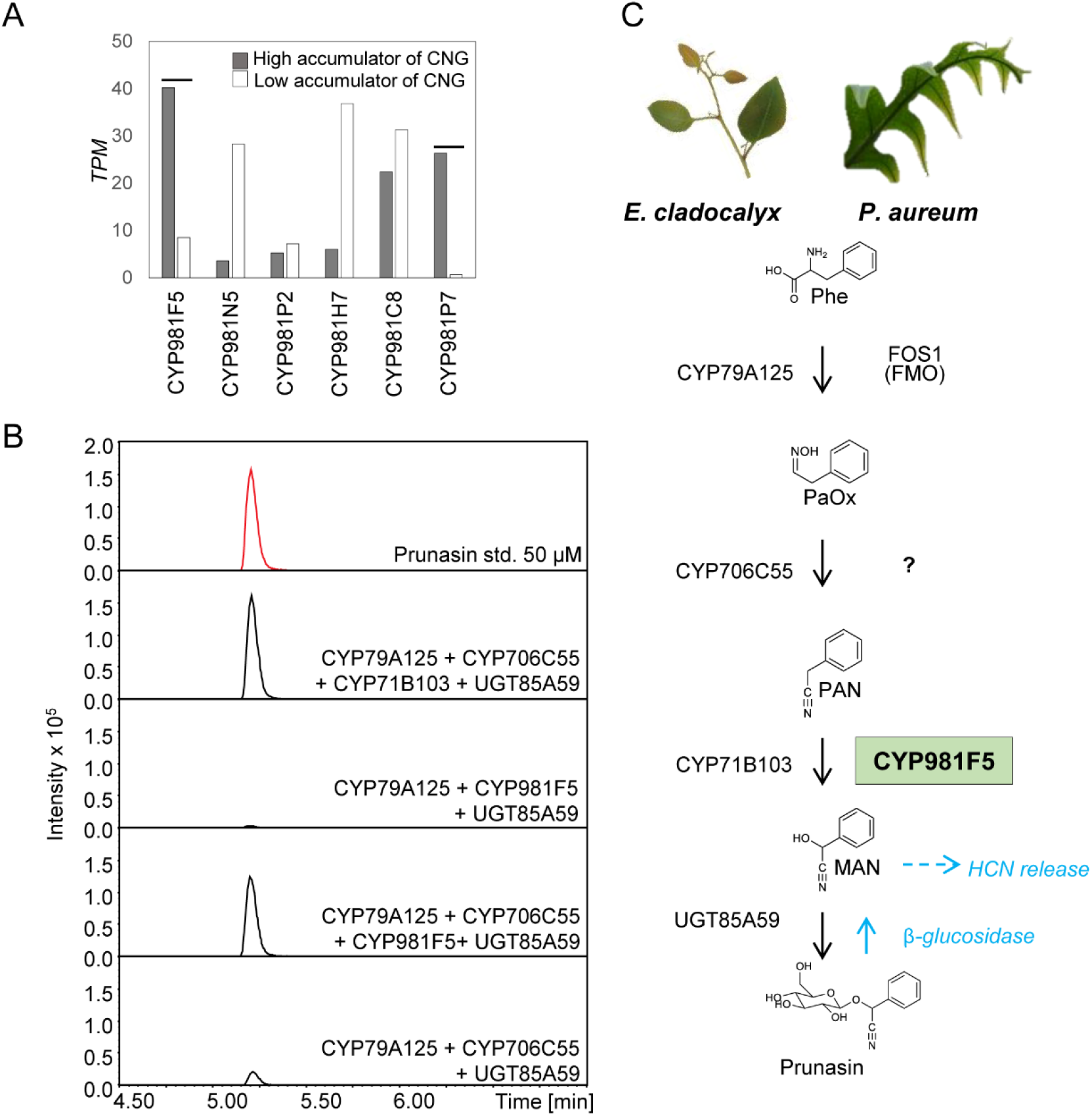
Functional characterization of PaCYP981F5. A) Expression profiles of CYP981s in P. aureum. Dark and white bars indicate transcript per million (TPM) from high and low cyanogenic glucoside producing tissues, respectively. Two CYP981 candidates were selected for biochemical investigations: PaCYP981F5 and PaCYP981P7. B) Extracted ion chromatograms of leaf extracts from N. benthamiana leaves transiently co-expressing PaCYP981F5 in combination with E. cladocalyx CYP79A125, CYP706C55, CYP71B103 and UGT85A59. C) Biosynthetic pathways for the cyanogenic glucoside prunasin from P. aureum and E. cladocalyx. Phe; Phenylalanine, PaOX; phenylacetaldoxime, PAN; Phenylacetonitrile, MAN; Mandelonitrile; CNG; Cyanogenic glycoside

Interestingly, *PaCYP981F5* was among the top 50 differentially expressed CYP genes in high- and low-cyanogenic glycoside accumulating tissues in *P. aureum* (Fig. S2). Looking beyond the CYP981 family, other highly differentially expressed *CYP*s belonged to more conserved CYP families such as CYP51, CYP74, CYP97 and CYP98 (Fig. S3/2). More likely, the differentiated expression levels of these could possibly reflect the different developmental stages of the two tissues.

*PaCYP981F5* and *PaCYP981P7* were chosen for functional characterization by transient expression in *Nicotiana benthamiana*, and co-expressed in different combinations with *Ec*CYP79A125, *Ec*CYP71B103 and *Ec*CYP706C55 catalyzing steps in prunasin biosynthesis in *E. cladocalyx* (Fig. 4B). This analytic system offered precise determination of the catalytic ability of the introduced *Pa*CYP981 enzyme. Metabolites were extracted from *N. benthamiana* leaves 5 days after agro-infiltration and analyzed using liquid chromatography tandem mass spectrometry (LC-MS/MS). As expected, co-expression of the four genes encoding the enzymes catalyzing prunasin biosynthesis in *E. cladycalyx* resulted in production of prunasin (Fig. 4B). Accumulation of a prunasin malonate ester conjugate was also observed (Fig. S3). Formation of malonylated derivatives has previously been observed upon heterologous production of prunasin and other natural products in *N. benthamiana* (Ting et al., 2013; Hansen et al., 2018; Thodberg et al., 2020) and is catalyzed by an endogenous acyltransferase in *N. benthamiana*. Malonylation may represent a detoxification response to the presence of prunasin and facilitate stacking of the prunasin molecules as inert storage stacks (Kallam et al., 2017). No prunasin was produced upon simultaneous replacement of *Ec*CYP706C55 and *Ec*CYP71B103 with *Pa*CYP981F5 (Fig. 4B), instead phenylacetaldoxime and phenylacetaldoxime-glucoside were observed to accumulate. In contrast, a simple direct substitution of *Ec*CYP71B103 with *Pa*CYP981F5 resulted in formation of a considerable amount of prunasin and the prunasin malonate ester. This finding demonstrates the ability of *Pa*CYP981F5 to hydroxylate phenylacetonitrile to form mandelonitrile. Trace amounts of prunasin were detected when the *Ec*CYP79A125, *Ec*CYP706C55 and *Ec*UGT85A59 were expressed, which is consistent with previous experiments (Hansen et al., 2018). *PaCYP981P7* did not show any activity in this experimental setup and base peak chromatograms did not reveal any metabolic differences from controls (Fig. S3).

### Identification of different cyanogenic glycosides throughout fern evolution

To assess the distribution of cyanogenesis across fern evolution, and compare whether the rise of this trait follows CYP981 family diversification, pinnae from 120 individual plants representing 94 species (Table S2) were collected and qualitatively screened for their ability to release hydrogen cyanide using the Feigl–Anger paper test (Feigl and Anger, 1966). Species were selected based upon their phylogeny with representatives being selected from 21 of the 37 evolutionarily defined clades. Nine cyanogenic species were identified, all belonging to the order Polypodiales (Fig. 3, S1). Of these, *Polypodium crassofolium* (Polypodiaceae) had not previously been identified as cyanogenic. *Stenochlae tenofolia* (Blechnaceaea), *Polypodium vulgare* and *Davallia canariensis* (Davalliaceae) only released hydrogen cyanide upon the addition of exogenous β-glucosidase. By adding exogenous prunasin, the species *Cythea cooperi* (Cyatheaceae), *Polypodium latum* and *Niphidium crassofolium* (both Polypodiaceae) were shown to have the capacity to hydrolyze prunasin as monitored by the release of hydrogen cyanide.

LC-MS/MS was used to determine the identity of the cyanogenic glycosides present in the nine cyanogen-containing species (Fig. 3). The mono-glucoside prunasin [(*R*)-mandelonitrile beta-D-glucopyranoside] accumulated in the species *S. tenofolia* (Blechnaceaea) and *Pteridium aquilinum* (Dennstaedtiaceae). The cyanogenic di-glycoside vicianin [(*R*)-mandelonitrile (6-*O*-alpha-L-arabinopyranosyl-beta-D-glucopyranoside)] accumulated in the remaining seven cyanogenic species *Phlebodium aureum, Polypodium vulgare, Polypodium crassofolium, Davalia thricomanoides, Davallia canariensis, Davallia ssp* and *Dryopteris filix mas*. In addition to vicianin, a second cyanogenic di-glycoside was also detected in the *Davallia spp*. (Fig. 5, S1,S4). The fragmentation pattern of this cyanogenic glycoside matched the literature data reported for lucumin [(*R*)-mandelonitrile (6-*O*-beta-*<u>D</u>*-xylopyranosyl-beta-D-glucopyranoside)] (Miller et al., 2006; Pičmanová et al., 2015).

**Figure 5.**
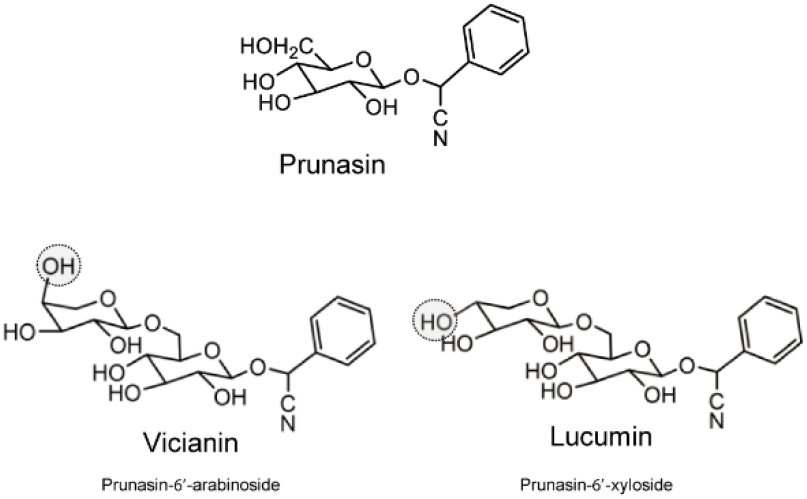
Cyanogenic glycosides in ferns. Cyanogenic glycosides identified by screening 105 fern species. Cyanogenic fern species harbor the phenylalanine-derived mono-glucoside prunasin and the corresponding di-glycosides vicianin and lucumin.

## Discussion

### A fern-centric view of plant CYPs

Phylogenetic analyses of the 8,000+ fern CYP sequences demonstrated both a molecular landscape unique to ferns, but also revealed an overall robust clan structure held throughout plant evolution. Ferns possess the highest number of CYP families compared to all other land plant lineages with 79 different families, 49 (60%) of which are fern-specific (Fig.1 and Fig. 2A). Analysis of CYP clans recognized deep-branching for the clans 51, 74, 97, 710, 711, and 727, as would be expected based on their strong conservation in the kingdom (Fig. 2). Despite this overall conserved structure, the inter-clan organization was sensitive to variance in alignment and input, especially concerning the position of the newly defined clans.

Many of the CYP families in ferns harbor a single member or a low member number as observed in the CYP98 and CYP51 clans. The CYP746 family is diminishing with evolutionary time and has disappeared in gymnosperms. CYP746 shows eclectic diversity and numbers in green algae, is present in mosses and liverworts but not in lycophytes, is sparse in ferns and extinct in seed plants. Thus, the lifetime of the CYP746 family seems to exhibit a spindle diagram shape, where the family begins at a point to widen to a maximum and then tapers off to zero over time (Gould et al., 1977).

Exploration into the OneKP dataset revealed a relatively high number of CYP73s and CYP74s per fern species (more than five copies per species). In comparison, CYP73s are typically present as a low number or single copy gene in gymnosperms and angiosperms. The CYP73 family catalyzes a conserved biosynthetic step in plants, functioning as a cinnamate-4-hydroxylase in the core phenylpropanoid pathway (Renault et al., 2017), whilst the CYP74 catalytic functions includes allene oxide synthase, hydroperoxide lyase or divinyl ether synthase (Laudert et al., 1996; Hughes et al., 2009). Some of the additional transcripts in ferns could derive from alternative splicing sites or as allele variants. The retention of higher copy numbers may also be explained by either neofunctionalization in pathways yet to be discovered, or by sub-functionalization as described for the CYP98 duplicates *CYP98A8* and *CYP98A9* in Brassicales (Liu et al., 2016). In the opposite way, the ” missing” CYP75 family in ferns, a family that hydroxylates the 3’- and 5’-position of the flavonoid B-ring (Holton et al., 1993), could be catalytically represented by the neighboring fern and lycophyte-specific CYP7332 and CYP7333 families (Fig 2B).

The fern CYPome provides insight into the molecular mechanisms in evolution that have been instrumental in orchestrating functional shifts or formation of new metabolic hubs. With a better understanding of the rise and fall of enzyme families, in comparison with the increased knowledge of metabolite profiles of specific species, deciphering of the chemo-evolutionary history of natural plant product formation becomes a possibility. More comprehensive and unbiased metabolite profiling in ferns, integrated with genomic/transcriptomic data, will guide the elucidation of currently unknown gene functions and/or biochemical pathways. The fern CYPome presented here provides an excellent platform for downstream identification of the molecular grids that have enabled plants (and ferns specifically) to evolve an immense number of unique metabolic pathways during evolution.

### The existence of ‘blooms’ of CYP families

The CYP981 family is the most abundant CYP family in ferns and is notably specific to this taxon. We mapped the occurrence of CYP981 subfamilies across fern phylogeny and their evolutionary relationships using the OneKP dataset (Table S1) and available genomic (Li et al., 2018) and in-house transcriptomic resources (Thodberg et al., 2020). The analysis showed that the evolutionary oldest CYP981 subfamily, CYP981D, was present in ancient ferns 370mya and displayed remarkable diversification and expansion along with the rise of the core leptosporangiate ferns 240 mya approximately. The presence of multiple CYPs very similar in their amino acid sequence (> 55% similarity) was common within individual fern species, and could represent recent gene duplication events. Further, the number of CYP subfamily members in a given fern species often differs even between close relatives. Such blooms, defined as recent lineage specific proliferation of paralogs, are not restricted to the CYP981 family, but are also observed for other CYP families in the fern CYPome (Table 1).

**Table 1:** Selected example of blooms from the plant kingdom.

| Fern blooms | Blooms in CYP71 family* |
| --- | --- |
| 23 CYP78 in *Azolla filiculoides* | 181 CYP71 in *Aegilops tauschii* (wheat genome D) |
| 30 CYP716B in *Azolla filiculoides* | 90 CYP71 in *Eucalyptus grandis* |
| 13 CYP716 in *Blechnum spicant* | 120 CYP71 in soybean, *Glycine max* |
| 14 CYP981AD in *Azolla filiculoides* | 122 CYP71 in rice, *Oryza sativa* |
| 29 CYP981 in *Pteridium aquilinum* | 347 CYP71 in tetraploid *Saccharum spontaneum* |
| 21 CYP990 in *Pteridium aquilinum* | 105 CYP71 in tomato, *Solanum lycopersicum* |
| 19 CYP993 in *Azolla filiculoides* | 223 CYP71 in potato, *Solanum tuberosum* |
| 13 CYP7331 in *Salvinia cucullata* | 442 CYP71 in hexaploid wheat, *Triticum aestivum* (3 genomes) |
| 12 CYP7339 in *Davallia fejeensis* |  |

\* derived from genomes and includes pseudogenes.

Gene duplications leading to blooms are stochastic events (Feyereisen, 2011; Dermauw et al., 2020). Subsequent selective pressure retains those of the new gene copies offering functional advantages to the organism. As sessile organisms, a key component enabling plants to adapt to and counteract all types of environmental challenges is the ability of plants to biosynthesize an immense array of natural products with diversified functionalities. Gene duplication and subsequent selection contributes to the contingent formation of natural products in separate lineages. CYPs are the key set of enzymes involved in the biosynthesis of natural products catalyzing coveted pathway steps. Given that ferns do not undergo a typical genome downsizing following whole genome duplication events as observed for other plant lineages (Liu et al., 2019) it is not surprising that ferns possess such high incidences of CYP family blooms. Furthermore, as this analysis relied mainly on transcriptomic resources, the number of fern CYP genes may be vastly underestimated.

The majority of extant fern diversity are found in the polypodiales and is the result of a second radiation initiated in the early-cretaceous period around 140 mya in parallel to the major uprising of angiosperms (Schneider et al., 2004). The dynamics of the CYP981 family and its potential role in production of bioactive natural products represent an effective response to the concurrent diversification of angiosperms and the parallel emergence of herbivores and pests. Thus, the CYP981 family retains a function analogous to the CYP71 family in higher plants.

Diversification of the CYP981 family in ferns may be considered equivalent to the large number of gene members in the multifunctional CYP71 family in seed plants (Fig. 2B). Ferns synthesize a diverse array of different natural products (Cooper-Driver et al., 1977; Alonso-Amelot, 2002; Kisielius et al., 2020) and it is highly likely that members of the CYP981 family catalyze some of the biosynthetic steps in these pathways. For example, the specific diversification of the CYP981 family in Pteridaceae, and the family-specific E, M and F CYP subfamilies, may be involved in the metabolic specialization of cytotoxic pterosins (Alonso-Amelot, 2002; Rasmussen, 2003; Lu et al., 2019). Further elucidation of natural product pathways in ferns shall clarify the degree of convergent evolution and neofunctionalization.

#### Does phenylalanine metabolism trigger CYP diversification?

The fern species *P. aureum* accumulates varying levels of cyanogenic glycosides in its different tissues (Thodberg et al., 2020) and formed the basis for a comparative transcriptomic approach to identify CYP candidates catalyzing the aldoxime to cyanohydrin conversion in cyanogenic glycoside biosynthesis. Specifically, we considered the six identified full-length members of the fern-specific CYP981 family. Two of these, *PaCYP981F5* and *PaCYP981P7*, were differentially expressed candidates with high expression in tissues accumulating high amounts of vicianin. Biochemical analysis showed that *Pa*CYP981F5 catalyzed hydroxylation of the expected intermediate phenylacetonitrile yielding the cyanohydrin mandelonitrile (Fig. 4B). Phenylacetaldoxime was not metabolized. This functionality supports the hypothesis on the recruitment of CYP981s in production of defensive natural products in response to environmental selective pressures.

The reaction catalyzed by *Pa*CYP981F5 is equivalent to the function of *Ec*CYP71B103 from *E. cladocalyx* (Hansen et al., 2018), and suggests that formation of mandelonitrile from phenylacetaldoxime similar to prunasin biosynthesis in *E. cladocalyx* requires a minimum of two distinct enzymes in *P. aureum*. This contrasts the biosynthetic route in other plant species where a single multifunctional CYP71 or CYP736 catalyzes the oxime to cyanohydrin conversion (Bak et al., 1998a; Jørgensen et al., 2011; Takos et al., 2011; Yamaguchi et al., 2014). These contrasting orchestrations of the cyanogenic glycoside pathway observed across the plant kingdom witnesses that the pathways evolved independently through combination of a series of enzymes with slightly different catalytic capacities. Notably, phenylalanine is the precursor for formation of the large class of natural products termed phenylpropanoids. Many of the products formed are bioactive or volatile, such as phenylethanol, phenylacetonitrile, phenylacetaldehyde, phenylacetic acid, phenylethylamine that plays roles in attracting pollinators or in defense against pathogenic organisms (Pichersky et al., 2006; Tieman et al., 2006; Schieber and Wüst, 2020). We hypothesize that prior to the establishment of complete enzymatic scaffolds synthesizing the entire route to cyanogenic glycoside formation, the recruitment of enzymes happened in a more sporadic manner, building single step reactions, each with selective benefits. Maybe it is therefore not a coincidence that the phenylalanine derived cyanogenic glycoside biosynthesis have multiple players, as it potentially serves as a metabolic hub to maximize the biological machinery. This is also exemplified in poplar (*Populus trichocarpa*), a non-cyanogenic tree displaying an ostentation of phenylalanine-derived volatiles in a crossroad of pathways (Kahn et al., 1999; Irmisch et al., 2013; Irmisch et al., 2014; Günther et al., 2019).

As observed in plants, the conversion of an aldoxime into its corresponding cyanohydrin in invertebrates can either be catalyzed by a single multifunctional enzyme or by two distinct enzymes. The Burnet moth (*Zygaena filipendulae*) produces the cyanogenic glucosides linamarin and lotaustralin by the action of CYP405A2, CYP332A3 and UGT33A1 (Jensen et al., 2011). The multifunctional catalytic activity of CYP405A2 resembles that of plant CYP79s and CYP332A3 converts the oxime to the corresponding cyanohydrin similar to the multifunctional cyanogenic CYP71s and CYP736 in angiosperms. In the millipede *Chamberlinius hualiensis*, CYP320B1 has been shown to hydroxylate phenylacetonitrile to yield mandelonitrile, i.e. a single catalytic step in contrast to CYP332A3 that catalyzes two steps (Yamaguchi et al., 2017). Thus, CYP enzymes capable of catalyzing the nitrile to cyanoghydrin conversion in cyanogenic glycoside synthesis have arisen by convergent evolution at several independent occasions.

### Cyanogenesis throughout fern evolution

To assess the distribution of cyanogenesis across fern evolution, and compare whether the rise of this trait follows CYP981 family diversification, pinnae from 120 individual plants representing 94 species (Table S2) were collected and qualitatively screened for their ability to release hydrogen cyanide using the Feigl–Anger paper test (Feigl and Anger, 1966).

Cyanogenic glycoside biosynthesis provides an excellent experimental system to study the evolution of a two-component chemical defense system across the plant kingdom. To specifically assess the trait across fern evolution we tested 21 out of the 37 described fern subfamilies (Schneider et al., 2004), including all the Polypodiales but the Olendraeceae. Our screen is the first to include the presence of exogenously added β-glucosidase to enable detection of cyanogenic glycosides in fern species devoid of the β-glucosidase activity required for hydrogen cyanide release. Of the 94 fern species tested, 11% possessed the ability to biosynthesize cyanogenic glycosides whereas only 6% (nine species) were cyanogenic. All cyanogenic species fall within the Polypodiales order, following the diversification of the CYP981 family (Fig 3). Given that the cyanogenic trait may be polymorphic (Schreiner et al., 1984; Gleadow and Woodrow, 2000; Gleadow et al., 2003), some of the species identified here as acyanogenic may contain cyanogenic individuals. The recorded percentage of cyanogenic fern species is in line with results from previous fern screens (Nicola L.Harper Gillian A. Cooper-Driver T. Swain, 1976). Across the plant kingdom, 4-5% of the species examined are reported to be cyanogenic (Lizotte et al., 1986; Wajant et al., 1995; Miller et al., 2004; Gleadow et al., 2008), while the proportion of cyanogenic gymnosperms is only 1-2% (Luck et al., 2017).

LC-MS analysis verified prunasin and vicianin as the two main cyanogenic constituents present in ferns and also revealed their presence in different quantitative and relative ratios (Fig. S1). In addition, we discovered the presence of an additional phenylalanine-derived cyanogenic diglycoside lucumin in a *Davallia* sp, thus expanding the cyanogenic glycoside profile in ferns. Lucumin contains a primeverose disaccharide moiety composed of glucose and xylose residues, and is the bitter constituent in the bark of species from the angiosperm family *Sapotaceae* (Eyjólfsson, 1971) and present in the rare Australian rainforest tree *Clerodendrum grayi* (*Lamiaceae*) (Miller et al., 2006).

Prunasin is the dominant cyanogenic glucoside in the older lineages Pteridaceae and Dennstaedtiaceae, whereas the diglycosides vicianin and lucumin were dominant in the species belonging to Polypodiaceae, Davalliaceae and Dryopteridaceae. The difference in glycosylation patterns observed in cyanogenic glycosides within the fern clade suggests that the UGTs catalyzing glycosylation of prunasin represent a later add-on to the pathway (Thodberg et al., 2018).

The mapping of cyanogenic glycoside occurrence onto fern phylogeny suggests that biosynthesis most likely not originated in a common ancestor to the Polypodiales order, but is a result of repeated evolution (Fig. 3). We suggest that the highly conserved FOS1 enzyme, that converts phenylalanine to phenylacetaldoxime in two species, *Pteridium aquilinum* (Pteridaceae) and *P. aureum* (Polypodiaceae) arose in an ancestral Polypodiales 140 mya (I P P G, 2016; Thodberg et al., 2020)(Thodberg et al., 2020). This production of the reactive oxime intermediate may have triggered repeated recruitment of downstream biosynthetic enzymes and would thereby mirror the situation observed in cyanogenic glycoside synthesis in angiosperms, where the oxime intermediate is metabolized by repeated recruitment of CYPs from different families within the 71 clan (Takos 2011, Hansen et al 2018).

### The enzymes missing in cyanogenic glycoside biosynthesis in ferns

An enzyme catalyzing conversion of phenylacetaldoxime to phenylacetonitrile and the two UGTs catalyzing glycosylation of mandelonitrile to produce prunasin and of prunasin to produce vicianin remain to be identified. The identification of such enzymes may profit from the availability of the assembled transcriptomic resource (Thodberg et al., 2020). Conversion of phenylacetaldoxime to phenylacetonitrile in prunasin biosynthesis in *E. cladocalyx* is catalyzed by *Ec*CYP706C55 and represents an unusual CYP catalyzed dehydration reaction (Hansen et al, 2018). In some non-cyanogenic plant species, CYP-catalyzed dehydration of aldoximes has also been reported. Two CYP71Bs from poplar and CYP71AT96 from giant knotweed (*Fallopia sachalinensis*) catalyze a similar reaction in the synthesis of herbivore-induced nitriles (Irmisch et al., 2014; Yamaguchi et al., 2016). Likewise, CYP71A13 from Arabidopsis converts indole acetaldoxime to indole-3-acetonitrile in camalexin biosynthesis (Nafisi et al., 2007). A CYP enzyme would be expected to catalyze the oxime to nitrile step in cyanogenic glycoside production in *P. aureum*. However, the recruitment of a flavin monooxygenase in ferns in contrast to CYP79s in angiosperms pinpoints the need for an unbiased broad-targeting search among differentially expressed enzymes.

## Material and methods

### Plant material

Pinnae samples of all tropical non-Scandinavian fern species used in the colorimetric Feigl– Anger based screen were obtained from ferns growing in glass houses at the University of Copenhagen Botanical Garden in Copenhagen. Pinnae were collected from fronds at comparable stages of ontogeny. Harvested tissue material was snap frozen in liquid nitrogen and stored at −80 °C until use. Scandinavian species (*Pteridium aquilinum, Dryopteris filix-mas*) where collected at Dronningens Bøge, Esrum lake, Nødebo (55°59’49.9” N 12°21’27.6” E).

### Hydrogen cyanide (HCN) potential and metabolite profiling of tropical ferns

Visual assessment of hydrogen cyanide potential was carried out based on the blue color staining of Feigl–Anger paper following disruption of fern tissues by a freeze–thaw cycle (Takos et al., 2011; Blomstedt et al., 2012). To avoid false negative results from cyanogenic glucoside containing fern tissues lacking the β-glucosidase, all plant tissues were also tested following addition of surplus amounts of a β-glucosidase extract from the *Lotus japonicus (cyd-1)* mutant, which does not produce cyanogenic glucosides (Takos et al., 2010; Blomstedt et al., 2012)

### LC-QTOF-MS analysis of cyanogenic glycoside content of ferns

Fern tissue (approx. 30 mg) was weighed and boiled in 85 % methanol (v/v, 300 μL) for 5 min. The vial was transferred to an ice bath and the material macerated with a small pestle. The supernatant obtained after centrifugation (13.000 x *g*, 1 min) was filtered (0.45 μm low-binding Durapore membrane) and diluted 1:5 in water prior to LC-MS analysis and quantitation of the prunasin content using a standard curve prepared from chemically synthesized pure prunasin (Møller et al., 2016) ([0.1 μM, 10 μM, 25 μM, 50 μM, 75 μM]. Single concentration mixtures of other authentic chemically synthesized compounds were used: sambunigrin [50 μM], linamarin [50 μM], prunasin [20 μM] (Møller et al., 2016). A crude vicianin extract was obtained from grinded and methanolic extraction of *Vicia sativa* seeds.

LC-MS of extracts of infiltrated *N. benthamiana* leaves was carried out either as described above using a Dionex Ultimate 3000RS UHPLC (Thermo Fisher Scientific) system with DAD detector and fitted with a Phenomenex Kinetex® XB-C18 column (1.7 μm, 100 × 2.1 mm; Phenomenex, US) operated at 40°C and with a flow rate of 0.3 mL/min. The mobile phases were: (A) 0.1 % HCOOH (v/v) and 50 mM NaCl; (B) 0.1 % HCOOH in MeCN (v/v). The gradient program: 0–1 min, isocratic gradient 5 % B; 1–7 min, linear gradient 5 %–70 %; 7– 8 min, linear gradient 70–100 %; 8–10 min isocratic 100 % B; 10–11 min, linear gradient 100–5 % B; 11–16 min, isocratic 5 % B. The UHPLC system was coupled to a compact™ qToF (Bruker Daltonics) mass spectrometer operated in negative ESI mode from 50 to 1200 *m/z*. Raw data was processed using Compass DataAnalysis software (version 4.2, Bruker Daltonics).

### Identification and quantification of Vicianin

An 85% MeOH-extract of ground *Vicia sativa* seeds was used as a source of vicianin in the LC-MS and thin layer chromatography (TLC) experiments. The presence of vicianin in the elution profiles was determined based on its accurate mass and UV absorption, respectively. Quantification of vicianin was carried out using a dilution series of an amygdalin standard (Sigma-Aldrich/Merck), as this cyanogenic di-glucoside is structurally comparable to vicianin and expected to behave similarly with respect to their degree of ionization and UV absorption.

### Isolation, cloning and heterologous expression of PaCYP981 candidate genes

The two ORFs harboring the full-length sequence of *PaCYP981F5* and *PaCYP981P7* (most upstream methionine) without codon optimization was fitted with attb1 and attb2 Gateway cloning sites: attB1: ggggacaagtttgtacaaaaaagcaggct, attB2: ggggaccactttgtacaagaaagctgggt and synthesized by GenScript. The fragment was cloned into the *E. coli* plasmid vector pUC57. Each of the two constructs were sub-cloned by Gateway recombination from the pUC57 vector into the expression vector pJAM-1507. overnight cultures of *Agrobacterium tumefaciens* (AGL1) containing the expression constructs with the target gene sequence for either *PaCYP981F5* and *PaCYP981P7* under the control of CamV-35S promoter/terminator elements in pJAM1502 or the gene sequence for the gene-silencing inhibitor protein p19 (Voinnet et al., 2015) were harvested by centrifugation (4,000 x *g*, 10 min) and resuspended to OD_600_ 0.8 in water. After 1h incubation at ambient temperature, the *A. tumefaciens* cultures were used to co-infiltrate leaves of 3– 4 weeks old *N. benthamiana* plants. After 4–5 d, leaf discs (1 cm diameter) were excised from infiltrated leaves, frozen in liquid nitrogen and subsequently ground and extracted in 200 µl 85 % MeOH (v/v) for metabolite profiling as described above.

### Phylogenetic Analysis

Derived amino acid sequences from the *Phlebodium aureum* transcriptomes classified as CYP981 sequences and CYP amino acid sequences classifying to the CYP981 family (>40%) as obtained from transcriptomes at OneKP project (*ualberta*.*ca/oneKP)* or as published(Thodberg et al., 2020). The original set of fern sequences from Salvinia and Azolla was downloaded from Fernbase (www.fernbase.org) by keyword search for P450. Both databases were blast searched using fern query sequences until only minor pseudogene P450 fragments were found. Nucleotide sequences were translated using the Virtual Ribosome 2.0 (services.healthtech.dtu.dk). The protein sequences were batch blast searched against all named plant cytochrome P450 sequences using NCBI Standalone BLAST software and a confidential database. The sequences were assigned CYP names based on the blast identity with known sequences. The fern CYPome was constructed with 79 sequences (Fig. 2A). The phylogenetic analysis utilized the maximum-likelihood method following ClustalW alignment in MEGA8. A Neighbor-joining tree was computed from the protein sequences utilized in Fig 2.B using CLUSTAL Omega at EBI (ebi.ac.uk). Sequences are deposited at Data S2.

## Accession numbers

Sequence data from this article can be found in the EMBL/GenBank data libraries under accession numbers; CYP981F5, XX000000 (pending) and CYP981P7 XX000000 (pending).

## Author contribution

Conceptualization, S.T., D.R.N., A.M.T., B.L.M. and E.H.J.N.; Methodology, S.T., C.C.H., D.R.N, A.M.T., B.L.M. and E.H.J.N.; Investigation, data curation and analysis, S.T., C.C.H., A.M.T, M.P. D.R.N. and E.H.J.N.; Writing – Original Draft, S.T. and C.C.H.; Writing – Review & Editing, S.T., C.C.H., A.M.T.,, B.L.M., D.R.N. and E.H.J.N.,; Funding Acquisition, B.L.M. and E.H.J.N.

## Acknowledgments

The authors gratefully acknowledge the Botanical gardens in Copenhagen and Århus, Denmark for guidance and access to plant materials. We would like to thank Amalie Kofoed Bendtsen, Mette Sørensen and the student group from 2018 for aiding the fern collection, and Rebecca E Miller for providing the lucumin extract from *Clerodendrum grayi*. Further, we are very thankful to gain early access into the 1000 plant transcriptome consortium, OneKP catalyzed by Carl J. Rothfels (University of California, Berkeley). This work was supported by the VILLUM Center for Plant Plasticity (VKR023054) (B.L.M.); the European Research Council Advanced Grant (ERC-2012-ADG_20120314) (B.L.M.); VILLUM Young Investigator Grant (VKR013167) (E.H.J.N.); a Danish Independent Research Council Sapere Aude Research Talent Post-Doctoral Stipend (6111-00379B) (E.H.J.N.); and a Novo Nordisk Emerging Investigator Grant (Grant No. 0054890) (E.H.J.N.). The financial support is gratefully acknowledged.

