## Supplemental tables and figures for "The fern CYPome: Fern-specific cytochrome P450 family involved in convergent evolution of chemical defense"

### Supplementary Table 1

| SAMPLE | CLADE | ORDER | FAMILY | SPECIES | TISSUE TYPE |
|---|---|---|---|---|---|
| CAPN | Eusporangiate Monilophytes | Equisetales | Equisetaceae | *Equisetum diffusum* | developing shoots |
| JVSZ | Eusporangiate Monilophytes | Equisetales | Equisetaceae | *Equisetum hyemale* | sterile leaves/branches |
| NHCM | Eusporangiate Monilophytes | Marattiales | Marattiaceae | *Angiopteris evecta* | developing shoots |
| UXCS | Eusporangiate Monilophytes | Marattiales | Marattiaceae | *Marattia sp.* | leaf |
| BEGM | Eusporangiate Monilophytes | Ophioglossales | Ophioglossaceae | *Botrypus virginianus* | Young sterile leaf tissue |
| WTJG | Eusporangiate Monilophytes | Ophioglossales | Ophioglossaceae | *Ophioglossum petiolatum* | leaves, stalk, sporangia |
| QHVS | Eusporangiate Monilophytes | Ophioglossales | Ophioglossaceae | *Ophioglossum vulgatum* | |
| EEAQ | Eusporangiate Monilophytes | Ophioglossales | Ophioglossaceae | *Sceptridium dissectum* | sterile leaf |
| QVMR | Eusporangiate Monilophytes | Psilotales | Psilotaceae | *Psilotum nudum* | developing shoots |
| ALVQ | Eusporangiate Monilophytes | Psilotales | Psilotaceae | *Tmesipteris parva* | Young fronds |
| PNZO | | Cyatheales | Culcitaceae | *Culcita macrocarpa* | young leaves |
| GANB | Leptosporangiate Monilophytes | Cyatheales | Cyatheaceae | *Cyathea (Alsophila) spinulosa* | leaves |
| EWXK | | Cyatheales | Thyrsopteridaceae | *Thyrsopteris elegans* | young leaves |
| XDVM | | Gleicheniales | Gleicheniaceae | *Sticherus lobatus* | young fronds |
| MEKP | | Gleicheniales | Dipteridaceae | *Dipteris conjugata* | young leaves |
| TWFZ | | Hymenophyllales | Hymenophyllaceae | *Crepidomanes venosum* | young fronds |
| QIAD | | Hymenophyllales | Hymenophyllaceae | *Hymenophyllum bivalve* | young fronds |
| TRPJ | | Hymenophyllales | Hymenophyllaceae | *Hymenophyllum cupressiforme* | young fronds and sori |
| BIVQ | | Osmundales | Osmundaceae | *Osmundastrum cinnamomeum* | |
| YKSS | | Osmundales | Osmundaceae | *Osmunda regalis* | leaf |
| UOMY | | Osmundales | Osmundaceae | *Osmunda sp.* | gametophyte |
| VIBO | | Osmundales | Osmundaceae | *Osmunda javanica* | mature leaves |
| UWOD | | Plagiogyriales | Plagiogyriaceae | *Plagiogyria japonica* | young leaves |
| IXLH | | Polypodiales | Polypodiaceae | *Polypodium hesperium* | young sterile leaves - B type |
| GYFU | | Polypodiales | Polypodiaceae | *Polypodium hesperium* | young sterile leaves - A type |
| YLJA | | Polypodiales | Polypodiaceae | *Polypodium amorphum* | young sterile leaves |
| CJNT | | Polypodiales | Polypodiaceae | *Polypodium glycyrrhiza* | leaf (sterile?) |
| ZQYU | | Polypodiales | Polypodiaceae | *Phlebodium pseudoaureum* | leaf |
| UJWU | | Polypodiales | Polypodiaceae | *Pleopeltis polypodioides* | dehydrating fronds |
| ORJE | | Polypodiales | Polypodiaceae | *Phymatosorus grossus* | leaf |
| OQWW | | Polypodiales | Davalliaceae | *Davallia fejeensis* | leaf |
| NWWI | | Polypodiales | Lomariopsidaceae | *Nephrolepis exaltata* | young leaf |
| JBLI | | Polypodiales | Elaphoglossaceae | *Bolbitis repanda* | leaf |
| FQGQ | | Polypodiales | Dryopteridaceae | *Polystichum acrostichoides* | Mix of sterile and young fertile leaves. |
| RFRB | | Polypodiales | Hypodematiaceae | *Didymochlaena truncatula* | young fronds |
| WGTU | | Polypodiales | Hypodematiaceae | *Leucostegia immersa* | just unfurling leaf plus young leaf |
| PSKY | | Polypodiales | Aspleniaceae | *Asplenium nidus* | leaf |
| KJZG | | Polypodiales | Aspleniaceae | *Asplenium platyneuron* | leaves |
| OCZL | | Polypodiales | Diplaziopsidaceae | *Homalosorus pycnocarpos* | sterile leaves (maybe a few sporangia) |
| MROH | | Polypodiales | Thelypteridaceae | *Thelypteris acuminata* | young leaf |
| YJJY | | Polypodiales | Woodsiaceae | *Woodsia scopulina* | young sterile leaves |
| YQEC | | Polypodiales | Woodsiaceae | *Woodsia ilvensis* | young leaves |
| HTFH | | Polypodiales | Onocleaceae | *Onoclea sensibilis* | leaves |
| VITX | | Polypodiales | Blechnaceae | *Blechnum spicant* | young leaf |
| AFPO | | Polypodiales | Athyriaceae | *Athyrium sp.* | gametophyte |
| URCP | | Polypodiales | Athyriaceae | *Athyrium filix-femina* | young leaf |
| FCHS | | Polypodiales | Athyriaceae | *Deparia lobato-crenata* | fertile leaf |
| UFJN | | Polypodiales | Athyriaceae | *Diplazium wichurae* | fertile leaf |
| HNDZ | | Polypodiales | Cystopteridaceae | *Cystopteris utahensis* | Mostly sterile leaf tissue; possibly some fertile. |
| LHLE | | Polypodiales | Cystopteridaceae | *Cystopteris fragilis* | young leaf+unfurling fiddlehead; appeared sterile |
| XXHP | | Polypodiales | Cystopteridaceae | *Cystopteris fragilis* | fronds |
| RICC | | Polypodiales | Cystopteridaceae | *Cystopteris reevesiana* | leaf (perhaps with some developing sporangia) |
| YOWV | | Polypodiales | Cystopteridaceae | *Cystopteris protrusa* | sterile leaves, slightly older |
| HEGQ | | Polypodiales | Cystopteridaceae | *Gymnocarpium dryopteris* | young sterile leaves |
| MTGC | | Polypodiales | Dennstaedtiaceae | *Dennstaedtia davallioides* | |
| WCLG | | Polypodiales | Pteridaceae | *Adiantum aleuticum* | young sterile leaves |
| BMJR | | Polypodiales | Pteridaceae | *Adiantum raddianum* | leaf |
| DCDT | | Polypodiales | Pteridaceae | *Gaga arizonica* | sterile leaves, some very young |
| WQML | | Polypodiales | Pteridaceae | *Cryptogramma acrostichoides* | Mix of sterile and young fertile leaves. |
| ZXJO | | Polypodiales | Pteridaceae | *Parahemionitis cordata* | leaf |
| POPJ | | Polypodiales | Pteridaceae | *Pteris vittata* | fronds |
| FLTD | | Polypodiales | Pteridaceae | *Pteris ensiformis* | young leaves |
| UJTT | | Polypodiales | Pteridaceae | *Pityrogramma trifoliata* | leaf (perhaps with some developing sporangia) |
| YCKE | | Polypodiales | Pteridaceae | *Notholaena montieliae* | young leaves, possibly fertile |
| XDDT | | Polypodiales | Pteridaceae | *Argyrochosma nivea* | young leaves probably sterile |
| GSXD | | Polypodiales | Pteridaceae | *Myriopteris rufa* | young leaves, possibly fertile |
| SKYV | | Polypodiales | Pteridaceae | *Vittaria lineata* | leaf, mostly or entirely sterile (hard to tell) |
| NDUV | | Polypodiales | Pteridaceae | *Vittaria appalachiana* | gametophyte |
| NOKI | | Polypodiales | Lindsaeaceae | *Lindsaea linearis* | Young fronds and sori |
| YIXP | | Polypodiales | Lindsaeaceae | *Lindsaea microphylla* | Young fronds and sori |
| CVEG | | Salviniales | Salviniaceae | *Azolla cf. caroliniana* | sterile plants (leaves) |
| KIIX | | Salviniales | Marsileaceae | *Pilularia globulifera* | young leaves |
| CQPW | | Schizaeales | Anemiaceae | *Anemia tomentosa* | sterile leaf |
| PBUU | | Schizaeales | Lygodiaceae | *Lygodium japonicum* | mix of fertile and sterile leaf tissue |

Supplementary Table 2 *94 fern species screened for cyanogenic glycosides*

| | |
|---|---|
| *Acrostichum aureum* | *Microlepia speluncae* |
| *Adiantum capillus-veneris* | *Microsorum punctatum* |
| *Adiantum digitatum* | *Nephrolepis cordifolia* |
| *Adiantum formosum* | *Nephrolepis hirsutula* |
| *Adiantum hispidulum* | *Nephrolepis pectinata* |
| *Adiantum raddianum* | *Nephrolepis rivularis* |
| *Adiantum raddianum Var. raddianum* | *Niphidium crassifolium* |
| *Adiantum seemannii* | *Pallaea viridis* |
| *Aglaomorpha drynarioides* | *Pecluma* |
| *Aglaomorpha heraclea* | *Pellaea falcata* |
| *Amblovenatum oplentum* | *Phlebodium aureum* |
| *Angiopteris chauliodonta* | *Phlebodium latum* |
| *Angiopteris evacta* | *Phlebodium macaronesicum* |
| *Asplenium bulbiferum* | *Phlebodium scandens* |
| *Asplenium daucifolium* | *Phlebodium subauriculatum* |
| *Asplenium marinum* | *Phlebodium subauriculatum 'knightiae'* |
| *Asplenium nidus* | *Phlebodium vulgare* |
| *Asplenium scolopendrium 'angustifolium'* | *Phymatosorus pustulatus ssp. howensis* |
| *Blechnum brasiliense* | *Phymatosorus scolopendria* |
| *Blechnum gibbum* | *Platycerium bifurcatum* |
| *Blechnum moorei* | *Platycerium grande* |
| *Blechnum occidentale* | *Platycerium hillii* |
| *Blechnum occidentale ssp. minor* | *Platycerium wandae* |
| *Campyloneurum phyllitidis* | *Platycerium willinckii* |
| *Cyathea cooperi* | *Pleopeltis hirsutissima* |
| *Cyrtomium falcatum* | *Polypodium australe* |
| *Cyrtomium falcatum 'rochfordianum'* | *Psilotum nudum* |
| *Davallia felixmas* | *Pteridium aquilinum* |
| *Davallia* | *Pteris cretica* |
| *Davallia canariensis* | *Pteris cretica 'albolineata'* |
| *Davallia trichomanoides* | *Pteris ensiformis* |
| *Dennstaedtia globulifera* | *Pteris multifida* |
| *Dennstaedtia producta* | *Pteris quadriauritra* |
| *Dicksonia antarctica* | *Pteris tremula* |
| *Didymochlaena truncatala* | *Pteris vittata* |
| *Diplazium esculentum* | *Pyrrosia* |
| *Doodia caudata* | *Rumohra adiantiformis* |
| *Drynaria* | *Rumohra adiantiformis* |
| *Goniophlebium subauriculatum* | *Selaginella involvens* |
| *Hemionitis* | *Selaginella kraussiana* |
| *Hemionitis arifolia* | *Selaginella moellendorffii* |
| *Huperzia phlegmaria* | *Selaginella pulcherrima* |
| *Huperzia squarrosa* | *Stenochlaena tenuifolia* |
| *Lygodium japonicum* | *Tectaria cicutaria* |
| *Maranta Lev.* | *Tectaria gemmifera* |
| *Microgramma squamulosa* | *Tectaria incisa* |
| *Microlepia platyphylla* | *Woodwardia sp* |

Supplementary Figure S1

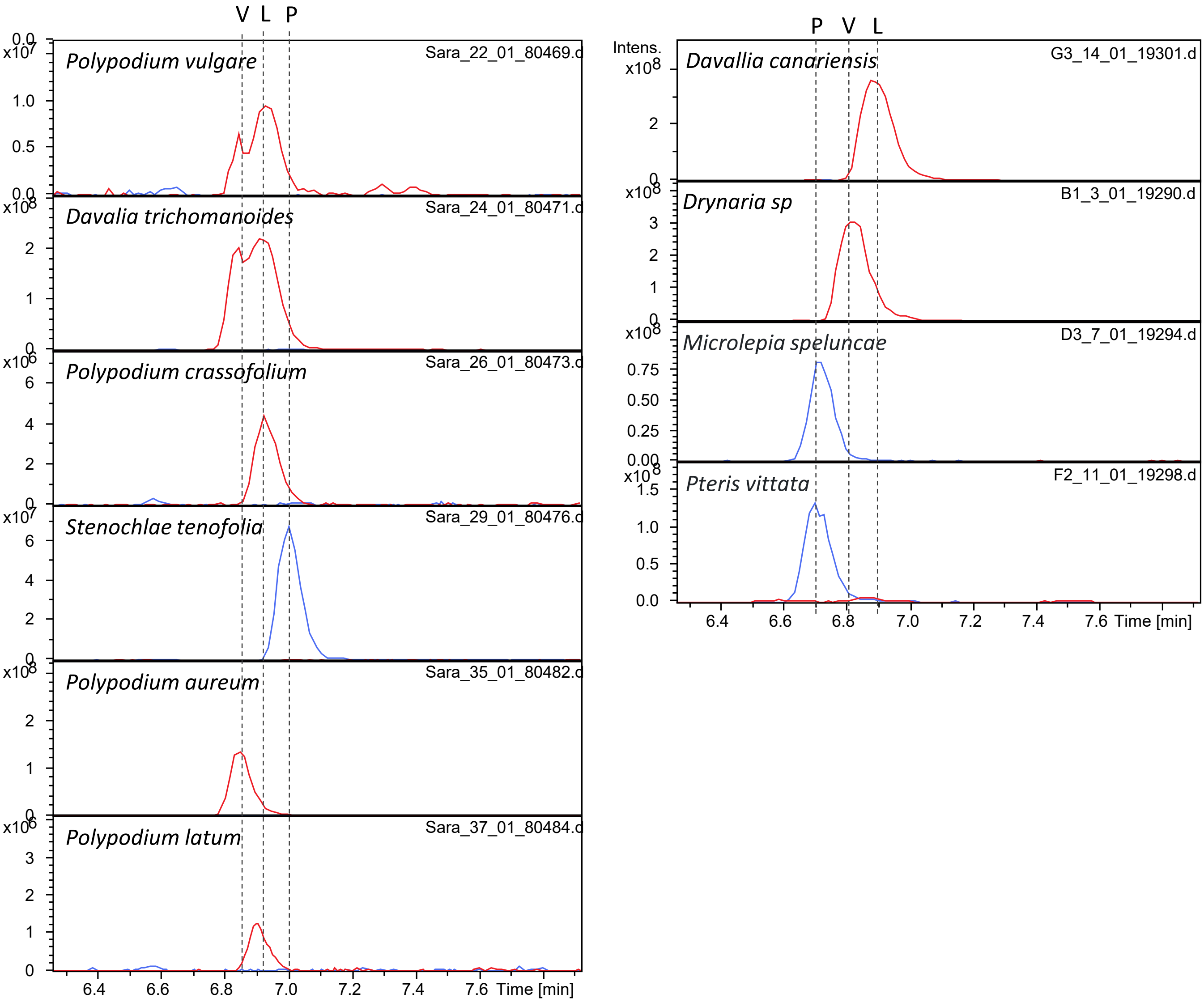

Supplementary Figure S2

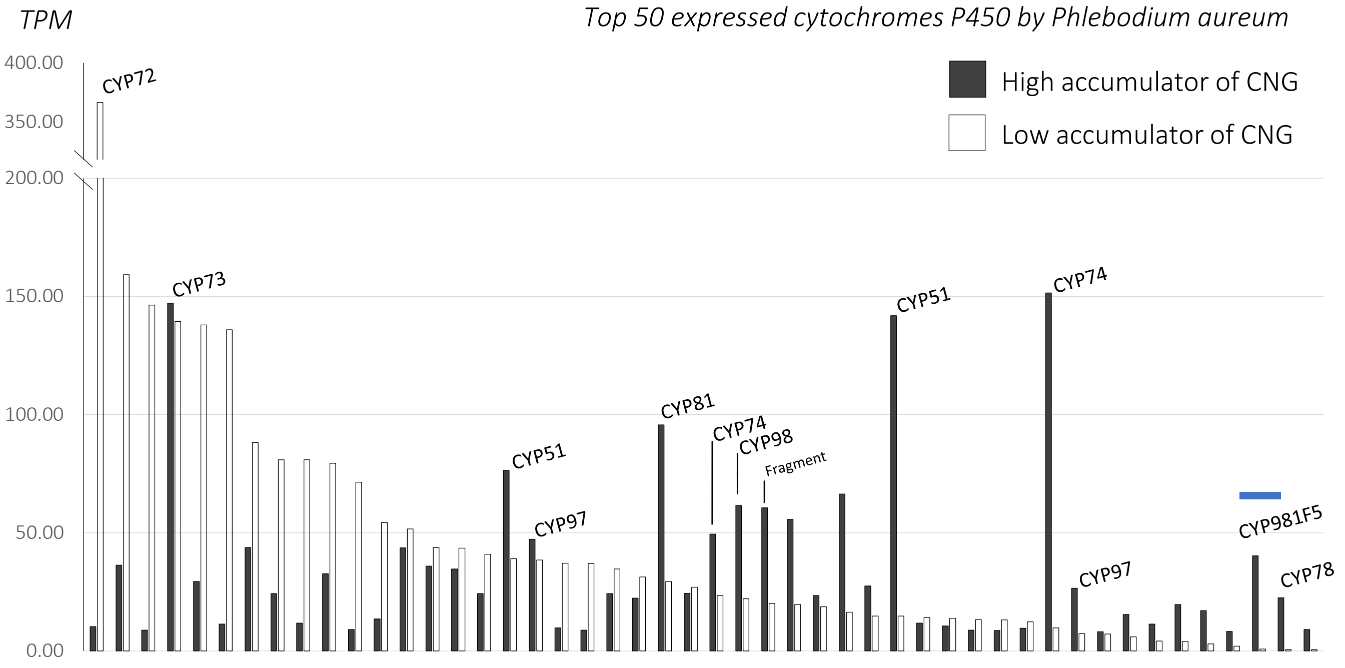


*Presentation of the 50 highest expressed contigs predicting cytochromes P450 in the high accumulator of cyanogenic glycoside in P. aureum. The contig are ordered according to the expression levels in the low cyanogenic glycoside containing tissue in order to highlight potential candidates.*

Supplementary Figure S3

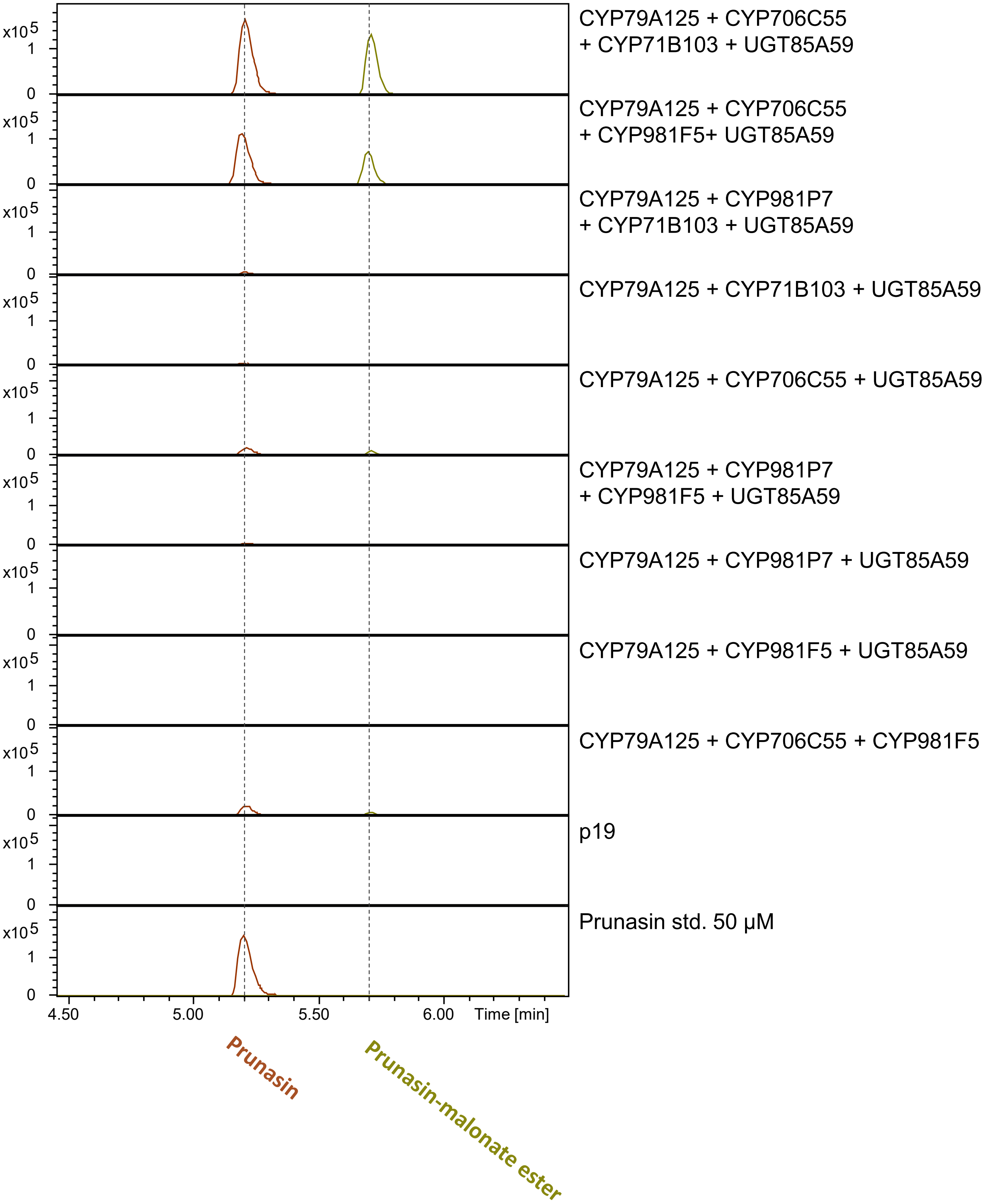

Supplementary Figure S4

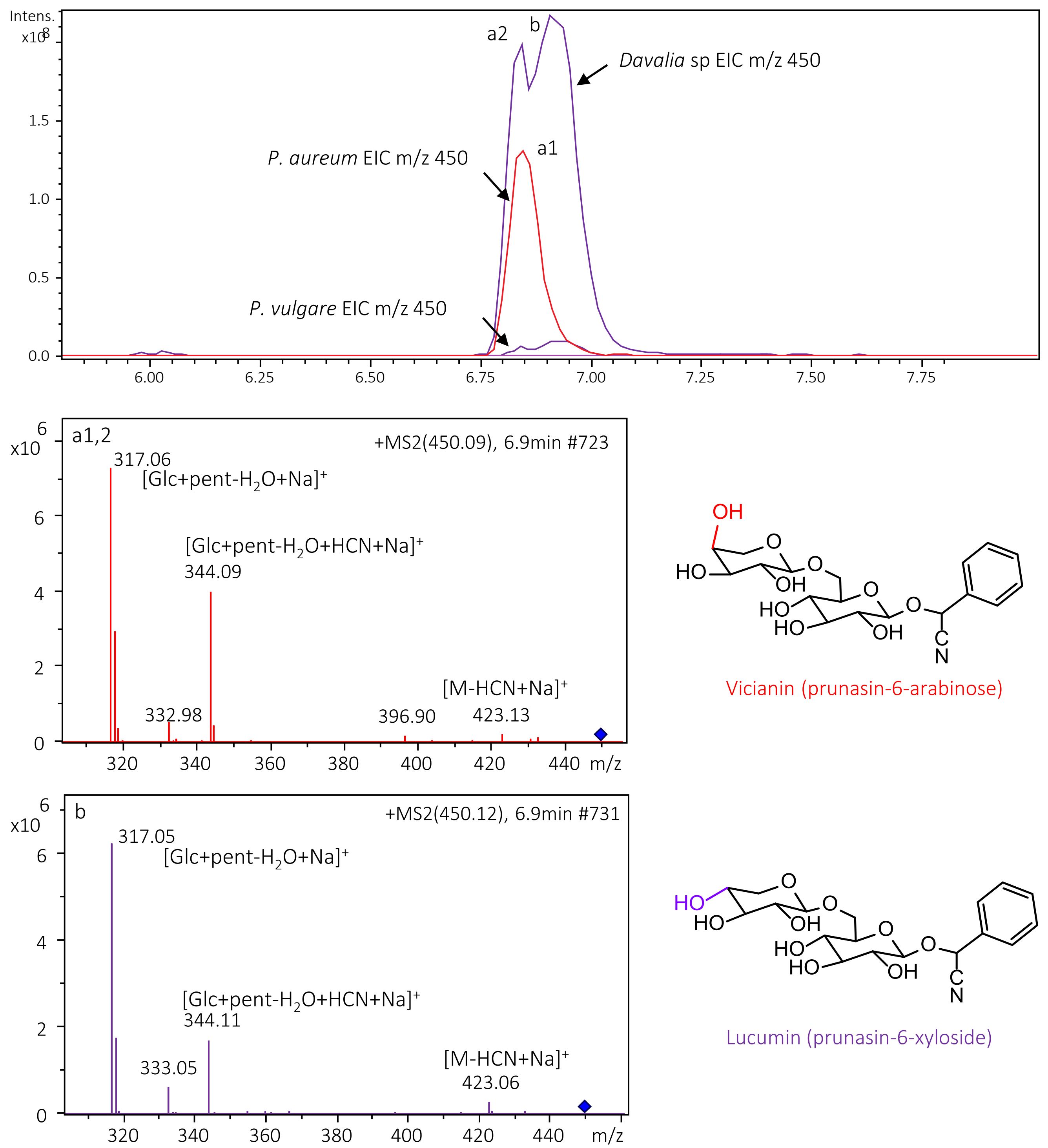
